# Identifying potential therapeutic targets for T-cell acute lymphoblastic leukemia using malignant networks and topological analysis

**DOI:** 10.1101/2025.10.08.681235

**Authors:** Rodrigo Henrique Ramos, Nicolas Carels, Rafaela Scardini, Letícia Lopes Franca, Adenilso Simao, Cynthia de Oliveira Lage Ferreira, Flávia Raquel Gonçalves Carneiro

## Abstract

T-cell acute lymphoblastic leukemia (T-ALL) is an aggressive and heterogeneous disease requiring new therapeutic targets. We identified overexpressed genes in T-ALL cell lines, built subnetworks using the IntAct interactome, and evaluated the topological role of each gene. An attack strategy, removing one gene at a time, was applied with eleven network measures and persistent homology. A positive control group of essential genes for T-ALL tumorigenesis was included. Clustering, largest connected component, and especially Betti 1 effectively distinguished control genes from others, showing that topological metrics are valuable for target identification. Notably, *NPM1* emerged as a key gene for maintaining network integrity, promoting proliferation, and ensuring survival, highlighting its potential as a promising therapeutic target in T-ALL.

**Significance of the study:** This study proposes an innovative approach to finding therapeutic targets for T-ALL based on oncogenic protein-protein interaction networks and the identification of the most effective analysis metrics following network attacks. These findings can be broadly applied to any neoplasm, seeking more personalized and efficient therapy.

## 1. Introduction

T-Cell Acute Lymphoblastic Leukemia (T-ALL) is one of the most aggressive forms of leukemia and occurs due to the malignant transformation of T-cell precursors. It is a genetically heterogeneous disease caused by the accumulation of lesions (genetic, molecular, or cellular abnormalities) that act in a multistep pathogenic process involving cell growth, proliferation, survival, and differentiation during the development of thymocytes. It represents approximately 15% and 25% of all Acute Lymphoblastic Leukemia (ALL) among the pediatric and adult cohorts, respectively [1].

With recent advances in therapy, 5-year overall survival rates have improved significantly, ranging from 74% to 91% in many contemporary pediatric clinical trials [2, 3]. The numbers are more modest if we consider adults separately, 30 to 60% [4, 5], although older adolescents and young adults treated with pediatric regimen had much better outcomes [6].

To reach these numbers, intensive therapy is required, causing many side effects. The frequent and multiple therapies can result in significant toxicity, contributing to both early morbidity and mortality, as well as serious long-term sequelae. Side effects include allergic reactions, gastrointestinal toxicity, myelosuppression, bone toxicity, increased risk of developing other tumors in the future, and neurological problems. Essential chemotherapy drugs used to treat ALL, such as L-asparaginase, methotrexate, vincristine, and nelarabine are neurotoxic and can affect 3.6–11% of children under treatment [7, 8, 9]. Moreover, within two years of diagnosis, 20% of patients with T-ALL will relapse, and the survival rate is less than 25% [10].

Therefore, a new therapeutic approach is necessary not only for relapsed patients but also to optimize the front line of treatment and minimize the short and long-term side effects of the current therapies. The heterogeneity of T-ALL lesions was confirmed through a broad analysis of 264 pediatric and young adult patients, including total sequencing, copy number analysis, and RNA sequencing [11]. They identified 106 target genes, half of which had not been previously described. More recently, Pölönen *et al*. [12] also performed an integrated analysis of genome and transcriptome sequencing of tumor and remission samples and identified recurrent sequence mutations for 16 previously unreported coding genes as well as non-coding genomic alterations. These studies indicate that potential new therapeutic targets still need to be explored.

In this report, we proposed a new method to identify therapeutic targets based on the construction of oncogenic protein-protein interactions (OncoPPIs) using overexpressed genes in Jurkat and MOLT-4 cell lines along with the IntAct Protein-Protein Interaction Network (PPIN) [13] to build the interactions. Hereafter, these OncoPPIs will be referred to as UP-OncoPPIs (Upregulated OncoPPIs) as only overexpressed genes were considered here.

It has been observed that the complexity of networks is correlated to patient survival [14, 15, 16] and tumor aggressiveness [17]. Furthermore, tumors present more complex protein interaction networks than normal tissues [18]. Understanding the OncoPPIs is fundamental for comprehending dysregulated pathways in tumors and for finding vulnerabilities related to interaction networks to guide precision therapeutic interventions [19, 20, 21].

Our UP-OncoPPIs provide a structural model for computational analysis, enabling more profound exploration of the functional relationships among genes involved in T-ALL. To evaluate the topological role of each gene within the networks, we employed the attack and impact strategy [22], a method frequently used in complex network studies. This approach measures how the removal of a vertex affects the overall structure and function of the modeled system. In our case, removing a vertex mimics a gene’s knockout. We used twelve metrics, including eleven conventional network science measures, and persistent homology (PH) as the twelfth measure to assess the impact of genes on higher-order structures within the network.

PH is a method of topological data analysis [23]. The main goal of PH is to determine interesting topological features of objects, such as connected components or higher-dimensional holes, considering different scales. Homology is commonly used to differentiate topological objects and counts the topological features in each dimension. In dimension 0, we denote by *b*_*0*_ the number of connected components. In dimension 1, *b*_*1*_ determines the number of one-dimensional holes. In general, *b*_*n*_ is called the *n*^th^ Betti number and determines the number of *n*-dimensional holes in a topological space. In cancer research, PH has recently been used to identify the importance of key genes in specific protein networks [24].

In this study, we included a positive control group of seven genes whose importance in T-ALL tumorigenesis has already been described or that have been evaluated in Jurkat and MOLT-4 lineages and are important for cell proliferation and survival [25, 26, 27, 28, 29, 30, 31]. These control genes, myelocytomatosis (*MYC*), heat shock protein family B (small) member 1 (*HSPB1*), serine/threonine kinase 1 (*AKT1)*, Y-box-binding protein 1 (*YBX1*), lysine (K)-specific demethylase 1A (*KDM1A*), heat shock protein 90 kDA alpha, class B, member 1 (*HSP90AB1*), and nucleophosmin 1 (*NPM1*), allowed us to assess which topological measures effectively distinguish them from other genes in the network, providing clues into which measures are necessary for characterizing therapeutic targets. This approach not only validated the relevance of certain topological features but also set the foundation for discovering new targets in leukemia.

It is important to note that *NPM1* is overexpressed in various hematologic and solid tumors [32, 33, 34] and is mutated in about 30% of acute myeloid leukemia patients [35]. Some studies have investigated the role of *NPM1* in T-ALL cells showing its overexpression in cell lines and its importance in cell viability [34, 31]. In this study we also analyzed the expression of *NPM1* and its role in cell viability and proliferation in the cell lines used as models for the UP-OncoPPIs, confirming some previous literature data and reinforcing the inclusion of this gene as part of the positive control group.

Our work proposes the use of UP-OncoPPIs as a structural model to better understand how gene inhibition affects malignant networks. By applying an attack strategy in which the removal of a vertex mimics a gene knockout, we aimed to assess the contribution of individual genes to network integrity through the analysis of a wide range of topological measures and persistent homology. We also incorporated a positive control group, which allowed us to assess the effectiveness of each metric in distinguishing key genes from others in the network. This strategy seeks to enhance the precision and reliability of approaches used to identify therapeutic targets.

## 2. Materials and Methods

### 2.1 In silico analysis

*RNA-seq sequences from T-ALL*: The bulk RNA-seq data (FASTq format) used in this report were from Jurkat (GSM2753018), and MOLT-4 (GSM2753021), two T-ALL derived malignant cell lines, as well as 12 samples of peripheral T lymphocytes from healthy donors (GSM1256828, GSM1256829, GSM1256830, GSM1256831, GSM1256832, GSM4144699, GSM4144700, GSM4144701, GSM4195996, GSM4195997, GSM4195998, GSM4195999). All the sequences were downloaded from the Gene Expression Omnibus (GEO) Dataset (https://www.ncbi.nlm.nih.gov/geo/) hosted on the National Center for Biotechnology Information (NCBI). More specifically, RNA-seq data from Jurkat and MOLT-4 were compared to the 12 control samples individually.

#### Data processing

After filtering out RNA-seq sequences for sequence stretch below Phred 33 using trimmomatic, we applied a series of Perl and Python customized scripts in the Galaxy environment [16] on BLASTn output to normalize and select up-regulated genes. Given that the frequency distribution of differentially regulated genes follows a symmetrical, Laplacian-like curve centered on zero of expression, we fitted a Gaussian distribution with 95% confidence and calculated the one-tailed critical value corresponding to a *p*-value of 0.025. The upregulated genes were considered as those whose expression was higher than the critical value.

#### Malignant networks

The human IntAct PPIN was preprocessed to retain only interactions with a probability greater than 50% of being true. Removing edges with low confidence scores is critical to ensuring reliable analyses [36]. To construct the UP-OncoPPIs’ networks for Jurkat and MOLT-4, we generated one consensus list of overexpressed genes for each cell line, considering there were 12 controls for comparisons. Therefore, genes that were found in at least 8 combinations were included to build the networks (Fig. 1A). The UP-OncoPPIs were then created by extracting induced sub-graphs from the IntAct PPIN, keeping only the largest connected component (LCC), and analyzed using the Python library NetworkX [37] with default settings. Finally, the subnetworks were visualized with the Gephi software [38].

**Fig. 1.**
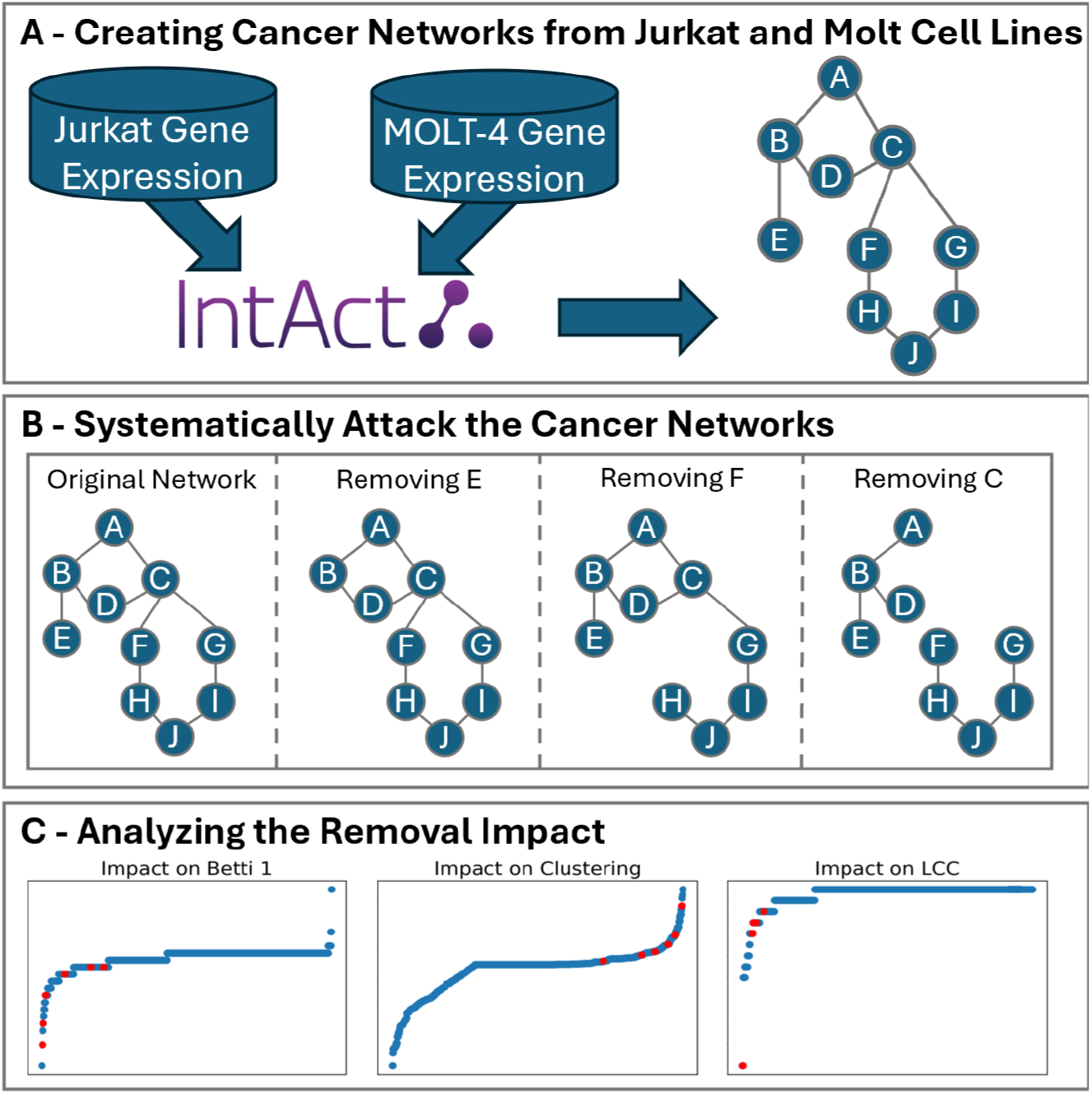
Method pipeline. Panel A illustrates the construction of UP-OncoPPIs from overexpressed genes in Jurkat and MOLT-4 cell lines. These genes were extracted from a preprocessed PPI to generate induced sub-graphs, forming the cancer networks. In panel B, network attacks were performed to systematically remove every gene in the network, one at a time, simulating the effects of gene knockout. In this example, the original network contains two Betti 1 structures: One formed by the vertices A, B, C, D and another by vertices C, F, G, H, I, J. The removal of vertex E does not affect any Betti 1 structure and is barely noticed by traditional measures. In contrast, the removal of vertex F breaks one Betti 1 structure. The removal of vertex C is the most impactful, destroying all Betti 1 structures and dramatically simplifying the network’s topology. In panel C, the topological impact of these attacks was analyzed, focusing on how they affect the structure and properties of the network, including changes in connectivity and higher-order features.

#### Positive control group

To evaluate the effectiveness of the applied measures in identifying relevant genes, we established a positive control group comprising overexpressed genes in Jurkat and MOLT-4, which played a key role in the proliferation and survival of these cells and/or were known to be crucial for T-ALL development. While other relevant target genes exist, our control group is limited to those within the network. The genes were *AKT1, HSP90AB1, HSPB1, KDM1A, MYC, NPM1*, and *YBX1*. The Jurkat network included all of them, while the MOLT-4 network did not contain *AKT1* and *HSPB1*.

#### Impact analysis of network attack

We removed one gene at a time from these malignant networks (Fig. 1B). This approach intends to mimic the biological process of gene silencing, effectively disrupting the protein interactions that the gene supports. By removing individual genes (vertices) along with their associated interactions (edges), we assessed network reorganization using topological measures, thereby elucidating the extent to which specific genes contribute to the structural integrity of the network.

The UP-OncoPPIs we analyzed are of moderate size, approximately 18 times smaller than the original PPIN from which they were derived but still have hundreds of vertices. To address this issue, we applied twelve network measures to assess the impact from different perspectives (Fig. 1C). The first six, LCC, density, entropy, modularity, degree assortativity and average path length are global network measures, whereas the subsequent five, betweenness centrality, closeness, eigenvector, clustering coefficient and degree represent average centralities metrics [39]. Although centrality measures characterize the importance of individual vertices, their average values offer insights into the network’s overall response to vertex removal. One can find a good explanation of each measure in [40].

Lastly, the twelfth measure, impact on Betti 1, explores higher-dimensional structures in the network, extending the analysis beyond vertices and edges [41, 42]. We assessed the impact of each vertex removal on the number of Betti 1 structures, related to persistent homology. Genes that impact the Betti 1 measure are engaged in more complex relational patterns, highlighting their significance in preserving the structural complexity of the network. This measure offers a deeper understanding of how a gene knockout or knockdown affects not only direct neighbors but also the broader network structure (Fig. 1B).

To assess the impact of the attack, we used *Mann-Whitney test*, which is a non-parametric statistical test used to compare differences between two independent groups when the data does not necessarily follow a normal distribution. In this analysis, data from both groups are ranked together, and the test evaluates whether there is a statistically significant difference between their distributions. The results include a U-statistic and an associated p-value, with significance determined based on a predefined alpha level (e.g., 0.05).

### 2.2 In vitro experiments

*Cells and cell culture:* MOLT-4 and Jurkat cell lines were obtained from the Cell Bank of Rio de Janeiro (Rio de Janeiro, Brazil, cat. #0176 and #0125, respectively). HEK293FT cell line for lentiviral production was kindly provided by Prof. Patricia Possik (Brazilian National Cancer Institute, INCA, Rio de Janeiro, Brazil). MOLT-4 and Jurkat cell lines were maintained in RPMI-1640 medium (Sigma) supplemented with 10% fetal bovine serum (FBS), L-glutamine, and penicillin-streptomycin. HEK293FT were cultured in Dulbecco’s modified medium (DMEM: Gibco) supplemented with 10% FBS, L-glutamine, and penicillin-streptomycin. Peripheral blood mononuclear cells (PBMCs) were obtained from healthy donors from INCA.

#### Lentiviral transduction

To generate *NPM1* knockdown cells, two independent shRNA sequences cloned into pLKO.1 vector were acquired (Sigma MISSION shRNA clones, NM_002520, TCRN0000062270 and TCRN0000062272). For control cells, the MISSION® pLKO.1-puro Non-Mammalian shRNA (SH002) vector was used. The viral packaging vectors pMDLg/pRRE, pHCMVg, and pRSVrev [43] were kindly provided by Prof. Patricia Possik (INCA, Rio de Janeiro, Brazil). To produce the lentiviruses, HEK293FT cells were transfected with the pLKO.1-shRNA (targeting *NPM1* or control) and the viral packaging vectors using lipofectamine 3000 (Thermo Fisher) according to the manufacturer’s instructions. Forty-eight hours after transfection, the supernatants containing the virus were harvested and used to infect T-ALL cell lines by centrifuging the cells for 45 min at 800 g and using 8 μg/ml polybrene. At 24 hours after infection, the medium was replenished with RPMI containing 10% FBS and 0.5 µg/ml puromycin to select the positive cells for 96 h. Thereafter, the cells were either plated for phenotypic assays or harvested and lysed for Western blot analysis.

#### Proliferation and MTT assay

Cell viability was evaluated by MTT (3-(4,5-Dimethylthiazol-2-yl)-2,5-Diphenyltetrazolium Bromide) assay at different time points (0, 24, 48, 72, and 96 hours). MOLT-4 and Jurkat cells transduced with shRNA specific for *NPM1* or control were plated in triplicate in 96-well plates at a density of 2.5 × 10^5^ cells/ml in 100 μl of RPMI. Ten microliters of MTT solution (5 mg/ml) were added to each well, followed by incubation for 4 hours at 37 °C. To solubilize the formazan crystals, 100 μl of a SDS-HCl solution (10% SDS, 0.01 M HCl) was added to each well, and the samples were incubated at 37°C for an additional 16 hours. Finally, the absorbance was measured at 570 nm. For proliferation, 10 μl of culture was taken from the same samples before the MTT addition and the trypan blue exclusion assay was performed.

#### Real-time quantitative PCR (qRT-PCR)

Total RNA was isolated from 5 × 10^6^ cells with RNeasy Mini Kit (Qiagen). 500 ng of RNA was reversed transcribed using the Transcriptase Reverse Superscript VILO Master Mix (ThermoFisher) according to the manufacturer’s protocol. cDNA was amplified using SYBR-Green PCR Master Mix (ThermoFisher), and *ACTB* was used as an internal control. The 2^−ΔΔCT^ method was applied to quantify the mRNA relative expression. The primers were: *ACTB*: (F) 5’ ACTCTTCCAGCCTTCCTTCC 3’, (R) 5’ TCTCCTTCTGCATCCTGTCG 3’ and *NPM1*: (F) 5’ CGGTTGTGAACTAAAGGCCGAC 3’ (R) 5’ CTCATCCTTTGCACCAGCCC 3’.

#### Western Blot

Control and treated cells were harvested and lysed in RIPA buffer (50 mM Tris pH 8, 150 mM NaCl, 1% de triton, 1% sodium deoxycholate, 0.1% SDS) with protease (Sigma-Aldrich) and phosphatase inhibitors (Sigma-Aldrich). The total lysate was quantified using a BCA protein assay kit (Pierce Biotechnology, Rockford, IL, USA) according to the manufacturer’s protocol. The protein samples (20 µg) were separated by SDS-PAGE and transferred to a Polyvinylidene fluoride (PVDF) membrane. Primary antibodies used are as follows: αNPM1 (AV34094, Sigma) and αACTIN (A5441, Sigma). The secondary antibodies were Goat anti-Rabbit IgG (31460, Thermo Fisher) and Goat anti-Mouse IgG (31430, ThermoFisher). The chemiluminescent signal was detected after the incubation of the membrane with the Super Signal West Pico PLUS Chemiluminescent Substract (3458, Thermo Fisher) in a Chemiluminescent Imaging System (Bio-Rad).

## 3. Results

### 3.1 Malignant networks

The IntAct PPIN was preprocessed to retain only interactions with a higher probability of being true. After this filtering, the interactome comprised 13,437 proteins (vertices) and 91,701 interactions (edges). RNA-seq data from Jurkat and MOLT-4 were compared to 12 samples of peripheral T lymphocytes from healthy donors. Genes that were upregulated in at least 8 of the 12 comparisons were selected to construct the networks. Therefore, we generated two consensus lists of overexpressed genes, one for Jurkat and another for MOLT-4 (Sup. Table 1), comprising 1418 and 1338 genes, respectively. Keeping only the largest connected component in the UP-OncoPPIs, the Jurkat network consisted of 731 vertices and 1,497 interactions (edges), while the MOLT-4 network contained 663 vertices and 1,230 interactions (Sup. Table 1 and Fig. 2). Notably, 58% of the vertices and 55% of the interactions in the MOLT-4 network were also present in the Jurkat network, while 53% of the Jurkat proteins and 45% of the interactions were found in the MOLT-4 network. Among the genes used to build the networks, we selected some that are essential for the cell lines’ proliferation and survival or are known to be critical for the development of T-ALL to compose the positive control group (Fig. 2).

**Fig. 2.**
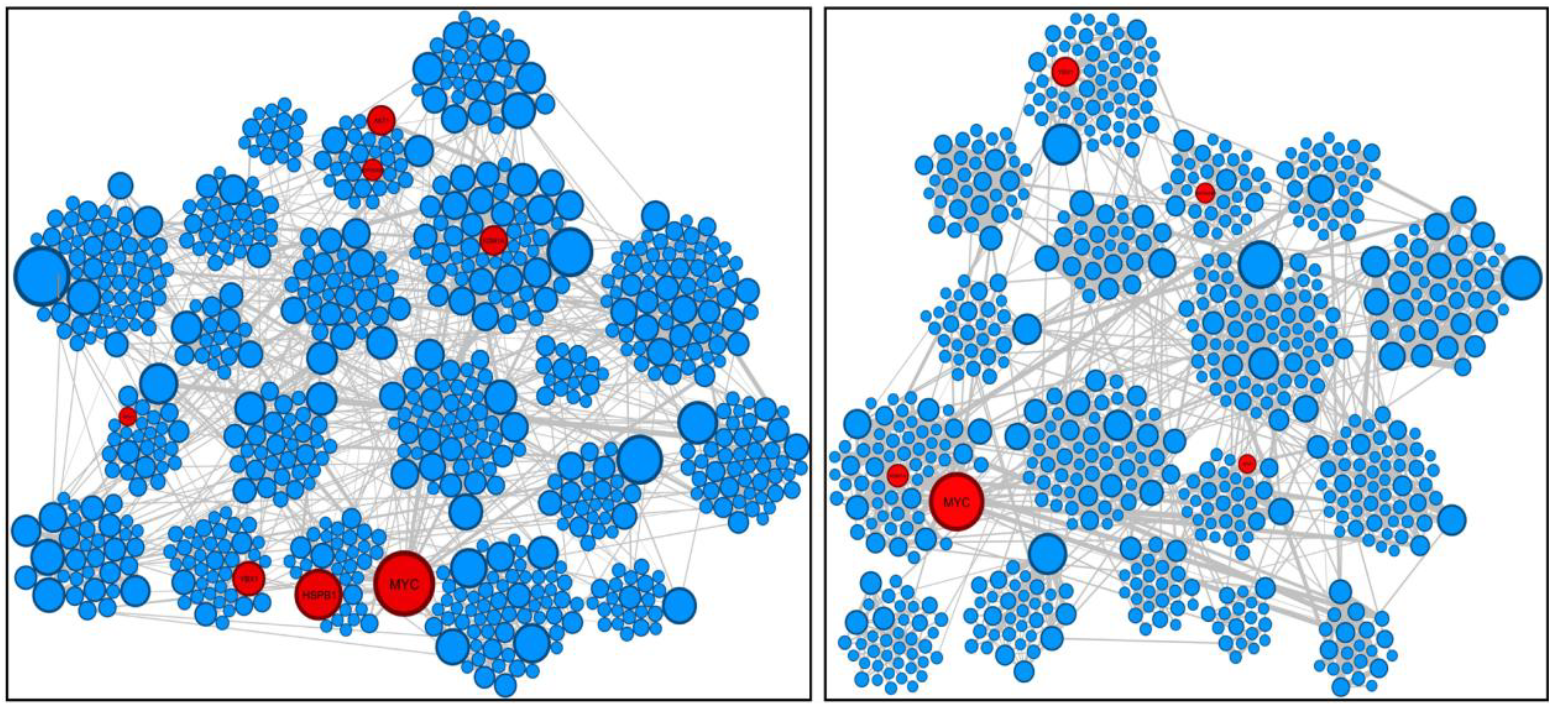
Jurkat (left) and MOLT-4 (right) networks. The plots emphasize communities and highlight the genes that composed the positive control group in red. The blue circles represent all the other genes in the network.

The network plots were generated using Gephi, with the vertices’ positions organized by communities. Vertex size is proportional to its degree (i.e., the number of interaction edges from the vertex considered), while edge thickness indicates the probability of the interaction being true. Interactions with a 75% or higher probability were predominantly located within communities. The control group genes were distributed throughout the network, with *AKT1* and *HSP90AB1* located within the same community in the Jurkat network (Fig. 2).

The Jurkat and MOLT-4 malignant networks exhibited average degrees of 4.1 and 3.7, respectively, with a median degree of 3. This distribution reflects the scale-free nature of the networks, characterized by numerous low-degree vertices and a few high-degree hubs. *MYC*, a well-established oncogene [44], is the vertex with the highest degree in both networks. *HSPB1*, present exclusively in the Jurkat network, has a high degree relative to the entire network and holds the highest degree within its community. *AKT1* (also exclusive to Jurkat) and *YBX1* function as community hubs, sharing their degree value and central role with several other genes. *HSP90AB1, KDM1A*, and particularly *NPM1*, exhibit typical degree values, both within their respective communities and in the broader network context (Sup.Table 2).

### 3.2 Attacking the Network

As previously described, we employed twelve topological measures to assess the impact of removing one gene at a time from the Jurkat and MOLT-4 networks. The resulting topological alterations induced by genes in the positive control group were then compared to those produced by the remaining genes in the networks (Fig. 3). An example to illustrate how topological measures may help to prioritize the selection of suitable targets for therapeutic purposes is presented in the first sub-plot of Fig. 3A, which shows the impact on Betti 1 in the Jurkat network. The complete network comprises 387 cycles. The removal of *HSPB1*, the first control group gene in the distribution, reduced the network to 371 cycles. Subsequently, the elimination of the second control gene, *MYC*, resulted in 373 remaining cycles, followed by *YBX1*, whose removal led to 378 cycles. In the Jurkat network, it is noticeable that *MYC* is positioned between two communities and maintains connections with others, indicating a high level of information flow (Fig. 2). When evaluating the impacts on modularity and betweenness (Fig. 3), the removal of *MYC* yielded the highest values in both metrics. This suggests that *MYC*’s removal triggers a network rewiring, increasing the number of communities and redistributing the information flow, previously concentrated on *MYC*, across other vertices, increasing the average betweenness. On the other hand, analyzing the Betti number in the MOLT-4 network reveals that the complete network comprises 219 cycles—168 fewer than in the Jurkat network (Fig. 3B, first plot). In this metric, *MYC* is the control group gene that produces the most significant impact. Similarly, in terms of betweenness centrality, *MYC* is also the most impactful gene, as also observed in the Jurkat network. However, for modularity in MOLT-4, *MYC* ranks as the second most impactful gene in the control group, surpassed by *YBX1*, which, in the Jurkat network, shares third place in this metric. These findings demonstrate that the two UP-OncoPPIs exhibit distinct topologies and that the same gene can have differing impacts depending on the network, reinforcing the importance of a comprehensive analysis that incorporates multiple structural measures. Although the absolute differences along the Y-axis are relatively small, given that we were assessing the impact of a single vertex removal at a time, distinct patterns emerged when analyzing the impact across all vertices. In some cases, the control group exhibited patterns that diverged markedly from that of other genes.

**Fig. 3.**
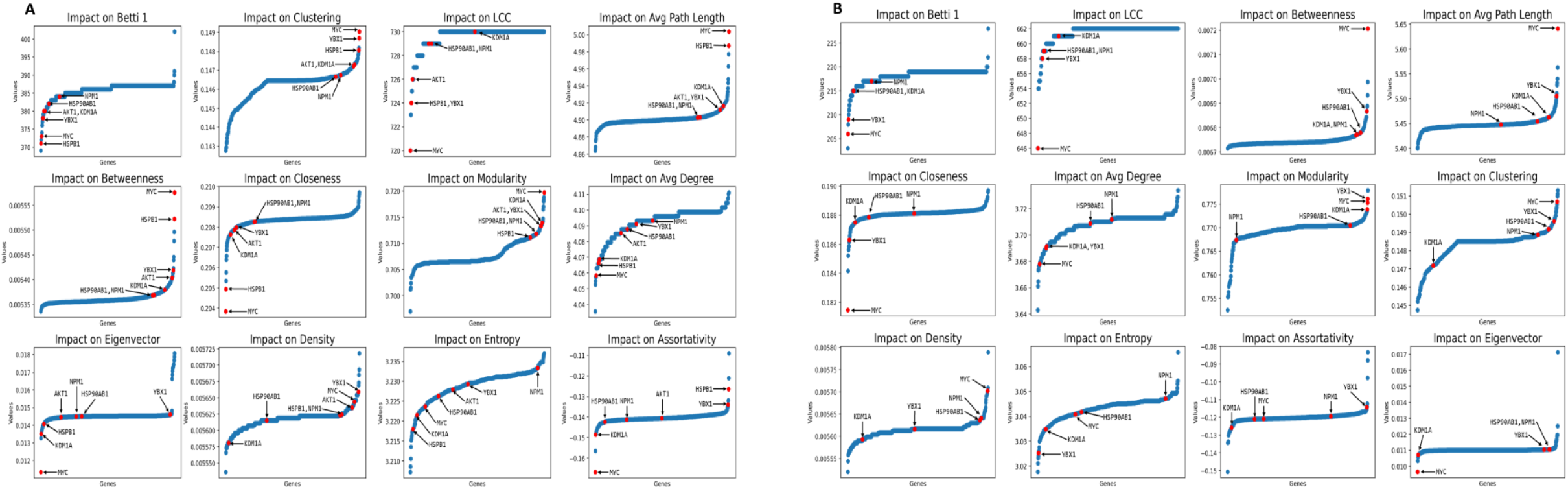
Impact distribution of all vertices using twelve measures on the Jurkat (A) and Molt-4 (B) malignant network. Red dots indicate genes of the positive control group, with their names pointed by arrows. Blue dots correspond to each remaining gene in the network (Sup. Table 2). The measures order follows the ranking of those that best separate the two groups. The Y-axis in each sub-plot represents the observed value after the removal of a gene for a given metric.

Upon analyzing both networks, we observed that certain metrics, such as assortativity and eigenvector centrality, exhibited nearly flat distributions. This suggests that, in most cases, the removal of individual genes does not substantially affect the network according to these specific measures (Fig. 3). In contrast, other metrics displayed more variable distributions, indicating that the impact of single-gene removal differs depending on the measure applied. Notably, in both networks, density and entropy demonstrated curved distributions; however, these metrics did not effectively differentiate control group genes from others. Conversely, measures such as Betti 1, LCC, and betweenness centrality revealed a tendency for control group genes to cluster at either the lower or upper extremes of the distributions. This pattern suggests that these metrics may hold potential for identifying candidate therapeutic targets.

To statistically assess the discriminative power of each metric, we applied the Mann-Whitney U test (Table 1). The metrics were ranked based on their ability to significantly distinguish control group genes from others in malignant networks. Metrics such as density, entropy, and assortativity yielded *p*-values above the 0.05 threshold and, therefore, were not considered relevant for characterizing the control group in either network. These results should be interpreted with caution and are used as a complement to the data derived from Fig. 3. It is important to note that the Mann-Whitney test evaluates ranked data and does not account for the magnitude of differences between values, rendering it less effective for metrics with low variance (i.e., flattened distributions). This limitation may explain why eigenvector centrality appears statistically significant for the Jurkat network but not for the MOLT-4 one, despite visual inspection suggesting that it does not meaningfully distinguish between the gene sets.

**Table 1.**
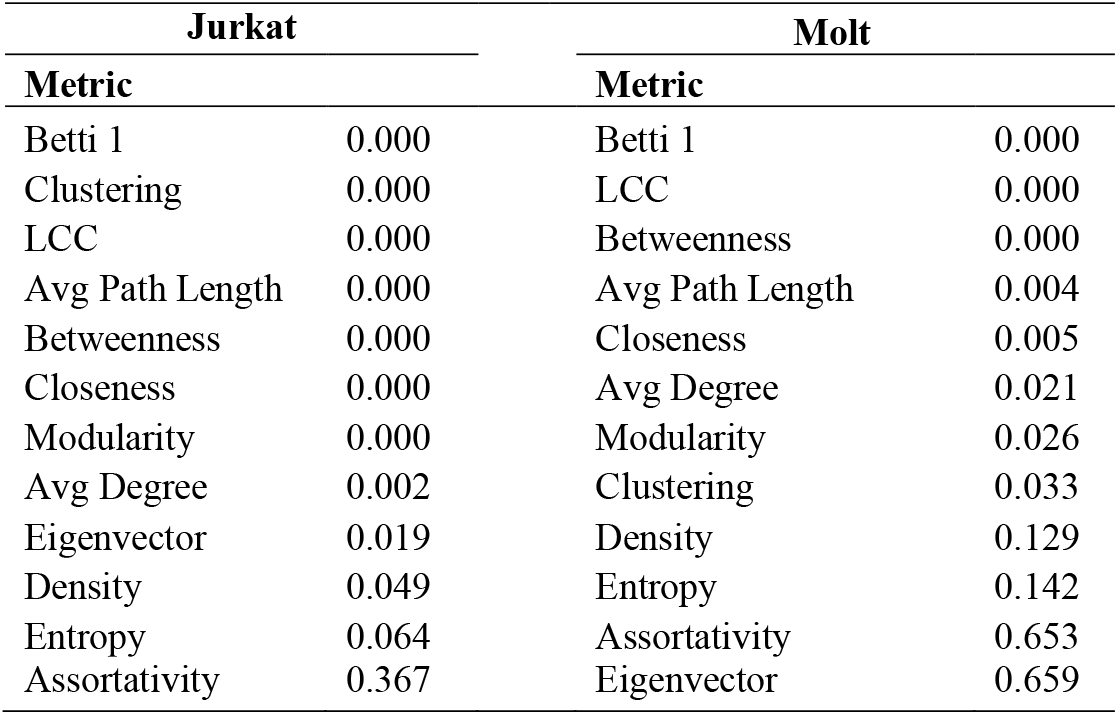
p-Values of Mann-Whitney test for network measures.

| Jurkat |  | Molt |  |
| --- | --- | --- | --- |
| Metric |  | Metric |  |
| Betti 1 | 0.000 | Betti 1 | 0.000 |
| Clustering | 0.000 | LCC | 0.000 |
| LCC | 0.000 | Betweenness | 0.000 |
| Avg Path Length | 0.000 | Avg Path Length | 0.004 |
| Betweenness | 0.000 | Closeness | 0.005 |
| Closeness | 0.000 | Avg Degree | 0.021 |
| Modularity | 0.000 | Modularity | 0.026 |
| Avg Degree | 0.002 | Clustering | 0.033 |
| Eigenvector | 0.019 | Density | 0.129 |
| Density | 0.049 | Entropy | 0.142 |
| Entropy | 0.064 | Assortativity | 0.653 |
| Assortativity | 0.367 | Eigenvector | 0.659 |

Following the network perturbation analysis, we selected the *NPM1* gene for *in vitro* validation, as it exhibited the least impact on the Betti 1 measure among the control genes. This implies that if *NPM1* significantly affects cell viability, the other control genes, associated with greater Betti 1 perturbations, are likely to demonstrate even stronger effects, thereby establishing *NPM1* as a conservative benchmark for therapeutic efficacy. Additionally, this gene was chosen due to its well-documented relevance in AML [35], and to further investigate its potential role in T-ALL within our model framework.

### 3.3 *NPM1* is overexpressed in T-ALL cell lines

The *NPM1* gene encodes the nucleophosmin protein, which has diverse functions, being involved in the correct centrosomes’ duplication, ribosome biogenesis, mRNA processing, chromatin remodeling, in addition to contributing to the maintenance of genomic stability and regulation of tumor suppressors [45]. In order to confirm the overexpression of *NPM1* in our model we performed RT-qPCR and western blot analysis comparing Jurkat and MOLT-4 cell lines with peripheral blood mononuclear cells (PBMC) from healthy donors. In both cases, *NPM1* was upregulated in the malignant cell lines (Fig. 4A and B).

**Fig. 4.**
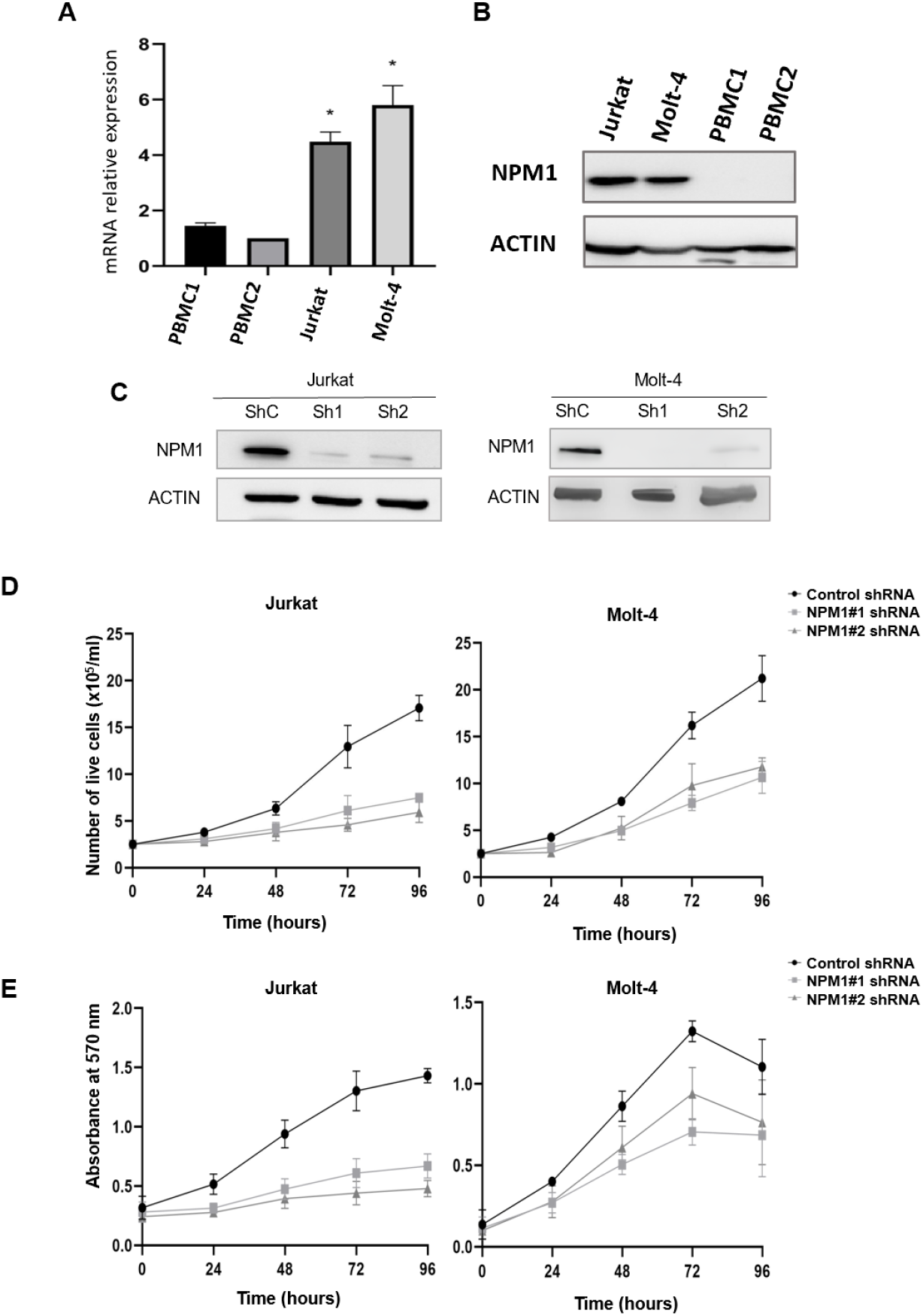
*NPM1* silencing reduces proliferation and viability in T-ALL cell lines. Overexpression of *NPM1* in Jurkat and MOLT-4 cell lines compared with PBMCs from healthy donors by qRT-PCR (A) and Western blot (B). Jurkat and MOLT-4 cell lines were transduced with shRNAs targeting NPM1. The protein levels of knockdown and control cells were detected by western blot (C). Cell proliferation was performed by trypan blue exclusion assay (D) and viability using the MTT assay (E). ShC: short hairpin Control; Sh1 and Sh2: Shot hairpins targeting *NPM1*. \**p* < 0.05

### 3.4 *NPM1* knockdown compromises T-ALL cell lines proliferation and viability

To understand the function of *NPM1* in our model, Jurkat and MOLT-4 cells were transduced with two independent shRNAs. Protein levels were significantly reduced in both cell lines (Fig. 4C) and knockdown cells showed a large decrease in cell proliferation (Fig. 4D) and viability (Fig. 4E) compared with control cells. These results demonstrated that *NPM1* may play a critical role in maintaining cell viability and proliferation in T-ALL cells.

## 4. Discussion

Significant efforts are currently being directed toward identifying target genes to support individualized treatment strategies for translational medicine. In this study, we analyzed 11 conventional network metrics, in addition to PH, to find new therapeutic targets for T-ALL and to identify the most effective metrics for this purpose. Furthermore, we established a positive control group composed of *MYC, YBX1, HSP90AB, HSPB1, AKT1, KDM1A*, and *NPM1*.

The role of *MYC* in hematological malignancies has been widely demonstrated [25, 46]. In T-ALL, *MYC* is important for tumor initiation and progression. Moreover, JQ1, a BET bromodomain 4 inhibitor, has been shown to reduce the levels of *MYC*, thereby decreasing the growth of human T-ALL cell lines as well as relapsed or treatment-resistant pediatric T-ALL cells *in vitro*. Treatment of Jurkat and MOLT-4 cell lines with JQ1 resulted in decreased *MYC* levels, impaired cell viability, and increased apoptosis [47]. Furthermore, the knockdown of *MYC* via shRNA in Jurkat led to reduced proliferation, while its inhibition through specific peptides decreased cell viability in MOLT-4 [48] [49]

*YBX1* is a multifunctional gene overexpressed in various human cancers and a member of the Cold Shock Domain family [50]. *YBX1* is overexpressed in Jurkat and MOLT-4 cell lines, as well as in cells derived from T-ALL patients. *YBX1* knockdown and knockout showed reduced cell viability, increased apoptosis, and G0/G1 phase arrest in both cell lines. Furthermore, *YBX1* depletion significantly reduced the leukemia burden in a human T-ALL xenograft and in a NOTCH1-induced T-ALL mice model in vivo [28].

*HSP90AB1* encodes a member of the heat shock protein (HSP) family and functions as a molecular chaperone. It is frequently overexpressed and contributes to tumorigenesis in many malignancies [51, 52]. Moreover, HSP90AB1 interacts with the intracellular domain of NOTCH1, regulating its stability and contributing to T-cell leukemogenesis [51]. More importantly, different HSP90 inhibitors reduced the viability and increased apoptosis in several T-ALL cell lines, including Jurkat and MOLT-4 [30, 53, 54].

*HSPB1* encodes a chaperone which is upregulated in many cancers and is associated with poor prognosis, therapeutic resistance, and tumor progression [55, 56]. HSPB1 inhibitors have been shown to increase Jurkat apoptosis, suggesting its role in this cell line survival [26]. We have selected *AKT1* gene for the control group since the PI3K/Akt/mTOR (PI3K) pathway has a crucial role in T-ALL and is activated in primary cells and several cell lines, including Jurkat and MOLT-4 [57, 27, 58]. The inhibition of AKT caused cell cycle arrest in G0/G1 phase, apoptosis, and reduction of viability in these cell lines [27] as well as increased sensitivity to asparaginase in primary T-ALL cells [57].

*KDM1A*, also known as *LSD1*, encodes a histone demethylase, which acts as an important gene expression modulator in eukaryotes. It is overexpressed in many solid and hematologic malignancies [59]. *KDM1A* silencing in MOLT-4 cells has been shown to reduce proliferation and induce apoptosis [29]. Other silenced T-ALL cell lines also showed impaired proliferation [60]. Moreover, *KDM1A* knockdown T-ALL cells failed to establish tumors in a mouse xenograft model [60].

Finally, we have chosen to analyze *NPM1* due to its well-established relevance in AML [33, 35] and to confirm that this gene can have a role in T-ALL tumorigenesis. In fact, we observed overexpression of *NPM1* in the cell lines used in this study, compared to PBMC samples. In addition, its knockdown resulted in impaired proliferation and survival. These findings underscore the need for further investigation of *NPM1* expression in T-ALL patients. The establishment of the control group with the aforementioned genes enabled the observation that integrating traditional network metrics with persistent homology improved the ability to identify key therapeutic targets in cancer networks. Some metrics worked better than others. Conventional measures, such as clustering and LCC, provided insights into local and global network connectivity patterns. For instance, genes exerting a significant impact on LCC were shown to play critical roles in maintaining network cohesion, emphasizing their structural importance in sustaining molecular interactions. Similarly, clustering highlighted crucial genes for local neighborhood integrity and community structure.

The inclusion of persistent homology, particularly Betti 1, added a refinement to the search for suitable therapeutic targets in oncology and in T-ALL, in particular. Unlike traditional metrics, which focus primarily on pairwise relationships and direct connectivity, Betti 1 captures higher-order topological features, thereby revealing the complexity of interactions within a network. However, the computational cost of persistent homology is exponential relative to both the size of the network and the dimensionality being analyzed [46]. This limitation restricts its application to larger networks or higher dimensional systems, underscoring the need for more efficient algorithms in future studies.

Our work differs from previous studies by employing a diverse set of topological measures that capture both local and global characteristics of network structure in order to characterize the control group. This multi-metric strategy enabled a detailed evaluation of how control group genes contribute to the network organization. The impact on the LCC suggests that control group genes are more critical than others in maintaining the network’s integrity, as their removal results in isolated components. The effects observed in the clustering coefficient and modularity indicate that the deletion of control group genes disrupts local connectivity and the structural organization of network communities. Furthermore, changes in betweenness centrality demonstrate that control group genes are pivotal for efficient information flow; their removal compels alternative pathways to assume this role, thereby increasing the average metric values for the whole network. However, among all the metrics analyzed, Betti 1 emerges as the most informative one for characterizing control group genes, being the only one that effectively differentiated this group from the others genes in the two cell lines studied. Consequently, genes exhibiting Betti 1 characteristics similar to those observed in the control group are likewise considered potential therapeutic targets.

## 5. Conclusion

This study integrates gene expression data, protein interaction networks, and topological analysis to investigate the topological features of therapeutic targets in leukemia, focusing on the Jurkat and MOLT-4 cell lines. By constructing malignant networks and systematically analyzing the impact of individual vertex (protein) removals, we aimed to identify topological measures that could effectively distinguish known therapeutic target genes from others. The results demonstrated that measures such as clustering, LCC, and, especially, Betti 1 effectively characterize the control group, with *p*-values significantly lower than those associated with other measures. These findings highlight the particular relevance of these topological measures in elucidating the structural significance of genes recognized as relevant in T-ALL and can be applied to other malignancies as well. Moreover, we reinforced the relevance of *NPM1* in cell proliferation and survival within our model. Therefore, the approach applied in this study may pave the way for discovering novel candidates for future therapeutic interventions.

AKT1: serine/threonine kinase 1
ALL: Lymphoblastic Leukemia
FASTq format: bulk RNA-seq
GEO: Gene Expression Omnibus
HSP90AB1: heat shock protein 90 kDA alpha, class B, member 1
HSPB1: heat shock protein family B (small) member1
KDM1A: (K)-specific demethylase 1A
LCC: Largest Connected Component
MTT: 3-(4,5-Dimethylthiazol-2-yl)-2,5-Diphenyltetrazolium Bromide
MYC: myelocytomatosis
NCBI: National Center for Biotechnology Information
NPM1: nucleophosmin 1
OncoPPIs: oncogenic protein-protein interactions
PBMCs: Peripheral blood mononuclear cells
PH: persistent homology
PPIN: Protein-Protein Interaction Network
T-ALL: T-cell acute lymphoblastic leukemia
UP-OncoPPIs: Upregulated OncoPPIs
YBX1: Y-box-binding protein 1

## Acknowledgements

This work received financial support from the Brazilian Federal Foundation for Support and Evaluation of Graduate Education (CAPES, Grant 88887.834984/2023-00), Inova FIOCRUZ Program (VPPCB-007-Fio18), Carlos Chagas Filho Foundation for Research Support of the State of Rio de Janeiro (FAPERJ: 211.325/2019 and E-26/211.404/2021). RHR was supported by the Federal Institute of Sao Paulo (IFSP). CLF and AS were supported by the University of Sao Paulo (USP). RSO was supported by fellowships from CNPq and INCA and LLS was supported by fellowships from INCA and FAPERJ. We are especially grateful to Dr. Patricia Possik (INCA) for kindly providing the lentiviral vectors and HEK293FT cells, and to Dr. João Viola and Dr. Maria do Socorro Pombo de Oliveira (INCA) for scientific discussions. We sincerely acknowledge the Program of Immunology and Tumor Biology (INCA) for providing access to their laboratory infrastructure and resources. We also extend our sincere appreciation to Amanda dos Santos Medeiros (INCA) for her technical expertise and support.

## Authors contributions

RHR and NC planned and executed experiments as well as wrote the manuscript, RS and LLS planned and executed experiments, CLF and AS co-supervised the study and wrote the manuscript, FRGC planned experiments, wrote the manuscript, contributed to the funding acquisition, and supervised this study.

## Ethical Approval

For the use of PBMCs, the study was approved by the Oswaldo Cruz Foundation (FIOCRUZ) and Brazilian National Cancer Institute (INCA) Ethics Committee under the registry numbers: CAAE 22005219.2.0000.5248 and CAAE 22005219.2.3001.5274, respectively. All the study methodologies conform to the standards set by the Declaration of Helsinki and experiments were undertaken with the understanding and written informed consent of all human subjects.

## Availability of data and materials

All code, input, and output files are on GitHub: https://github.com/RodrigoHenriqueRamos/Identifying-therapeutic-targets-for-Leukemia-using-malignant-networks-and-topological-analysis

---

**Supplementary Table 1 - Overexpressed vertices of Jurkat and MOLT-4 networks**. The spreadsheets indicate: Jurkat_Upregulated_12_Samples and MOLT4_Upregulated_12_Samples, which identify overexpressed genes by comparing the cell lines with 12 controls (peripheral T-cells from healthy donors) individually; Jurkat_GeneFrequency and MOLT4_GeneFrequency, which show the frequency at which each gene was upregulated across the 12 comparisons. The ones with a frequency higher than 8 were selected to generate the consensus list for each cell line; Jurkat_NetworkNodes and MOLT4_NetworkNodes comprised genes that formed the largest connected component of the UP-OncoPPIs and were used to build the final network for each cell line.

**Supplementary Table 2**. Connectivity degree of genes from the positive control group in the Jurkat and MOLT-4 networks.

**Supplementary Table 3**. All the network genes were ranked according to the impact of their removal, considering the different measures

