## Supplementary material for "Identifying potential therapeutic targets for T-cell acute lymphoblastic leukemia using malignant networks and topological analysis": Sup. Table 2

**Supplementary Table 2.** Connectivity degree of genes from the positive control group in the Jurkat and MOLT-4 networks.

|  | **Genes** | | | | | | |
| --- | --- | --- | --- | --- | --- | --- | --- |
| **Cell lines** | **MYC** | **HSPB1** | **YBX1** | **AKT1** | **KDM1A** | **HSP90AB1** | **NPM1** |
| **Jurkat** | 34 | 25 | 15 | 12 | 12 | 7 | 5 |
| **MOLT-4** | 37 | - | 15 | - | 10 | 8 | 7 |
